# Blue appendages and temperature acclimation increase survival during acute heat stress in the upside-down jellyfish, *Cassiopea xamachana*

**DOI:** 10.1101/2024.03.20.586014

**Authors:** Megan E. Maloney, Katherine M. Buckley, Marie E. Strader

## Abstract

Upside-down jellyfish (*Cassiopea sp.*) are highly tolerant to multiple abiotic stressors, including fluctuating temperatures associated with shallow marine habitats. This resilience may underlie the ability of *Cassiopea sp.* to inhabit a wide variety of tropical habitats across the globe. Additionally, *Cassiopea sp.* are marked by a conspicuous array of appendage coloration; individual medusae vary in the hue and number of oral appendages, which are often strikingly blue. The function of this coloration is not understood. We aimed to understand how extrinsic and intrinsic factors may shape thermal tolerance. Adult *Cassiopea xamachana* were collected from two sites that vary in daily temperature range within the Florida Keys and were subjected to acute lethal heat stress experiments. To quantify a whole-organism response to heat, we measured changes in bell pulsation, which likely plays a role in feeding, oxygen exchange, and symbiont uptake. Results show that *C. xamachana* from the two collection sites do not exhibit different responses to heat, suggesting that temperature fluctuations do not prime individuals for higher thermal tolerance. Additionally, *C. xamachana* with blue appendages survived significantly higher temperatures and exhibited less change in bell pulsation rates compared to non-blue individuals. Finally, color morphs were acclimated at either ambient (26 °C) or elevated (33 °C) temperatures. We found that acclimation at 33 °C, as well as appendage color in each treatment, led to higher survival under acute heat stress. Together, these findings highlight the importance of phenotypic plasticity and coloration in *Cassiopea* resilience during heat stress.

## Introduction

As global change continues to threaten marine ecosystems, rising sea surface temperature is the dominant stressor for many shallow water marine species (Filbee-Dexter et al., 2020; Lang et al., 2023; Mellin et al., 2019; Pörtner et al., 2019; Strydom et al., 2020). Ectotherms are particularly vulnerable to environmental variation, as even small temperature changes (1 – 2 ℃; Pinsky et al., 2019) can affect physiology and behavior (Lagerspetz & Vainio, 2006; Lang et al., 2023). In response to environmental change, ectotherms can adjust physiological traits (*e.g.*, respiration rate, metabolism, and reproduction; Guderley, 1990; Seebacher et al., 2015) through phenotypic plasticity (Ghalambor et al., 2007; Hartl & Conner, 2004; Jardeleza et al., 2022). This plasticity may enable some marine ectotherms to survive short-term heatwave events, which can help maintain genetic diversity for future selection to act upon (Botero et al., 2015; Gunderson & Stillman, 2015; Huey et al., 2012; Palumbi et al., 2014; Somero, 2010). Given the rapid pace of sea surface temperature increase, it is critical to quantify the role phenotypic plasticity plays in maintaining variation within and among populations of diverse marine species, and how this may contribute to organismal persistence in the face of environmental stress.

Plastic phenotypes are generally favored in the presence of predictable environmental variability (DeWitt et al., 1998; Goldstein & Ehrenreich, 2021; Moran, 1992), particularly when the costs associated with adjustment are minimal (Gavrilets & Scheiner, 1993; Pigliucci & Schlichting, 1998; Scheiner, 2018). As with most traits, plasticity itself can vary within and among populations, which may further affect the distribution of resulting phenotypes (Fuller et al., 2022). Natural experiments in coral reef systems have shown that individuals from environments characterized by high temperature variability, such as those experienced in back-reef pools and intertidal/shallow reefs, have enhanced thermal tolerance (Palumbi et al., 2014; Rivest et al., 2017; Safaie et al., 2018; Schoepf et al., 2022a). Increases in thermal tolerance as a result of plasticity may facilitate the persistence of individuals during the inevitable environmental change associated with climate change (Schaum et al., 2022).

Many marine invertebrates exhibit profound variation in color, which can be produced in one of several ways: structural coloration (*i.e.,* microstructures that selectively interfere with light; Parker, 1998), pigments and chromoproteins, or (Haddock & Dunn, 2015). In some marine invertebrates, color variation has been linked to physiological processes such as thermoregulation and stress response (Umbers, 2013). For example, green and red variants of the sea cucumber (*Apostichopus japonicus*) exhibit differential thermotolerance (Dong et al., 2010). In cnidarians, however, which exhibit particularly diverse color patterns from pigment-derived proteins (*e.g*., carotenoproteins) and fluorescent proteins (*e.g.*, green fluorescent protein [GFP]; Matz et al., 2002; Shagin et al., 2004), experimental evidence that supports a physiological role for coloration remains limited (Haddock & Dunn, 2015).

Studies investigating links between colors and ecological/evolutionary processes in marine invertebrates have primarily focused on the most common colors: reds, greens, and oranges (Muñoz-Miranda & Iñiguez-Moreno, 2023). In contrast, work on blue colors has been limited, largely due to the rarity of this color in marine habitats (Lawley et al., 2021; Newsome et al., 2014). One of the most striking examples of blue coloration in marine invertebrates is the upside-down jellyfish (*Cassiopea sp.*; (Y. Arai et al., 2017; Gamero-Mora et al., 2022; Medina et al., 2021; A. H. Ohdera et al., 2018). *Cassiopea xamachana* individuals exhibit a striking diversity of color variation, with many different patterns of deep blues and greens in oral appendages and throughout the mesoglea. This unique blue is produced by a chromoprotein known as Cassio Blue (Phelan et al., 2006), which has orthologs in several scyphozoan corals and jellyfish (Bulina et al., 2004; Lawley et al., 2021). Despite the high concentration of Cassio Blue in *C. xamachana* (up to 6% of total animal protein, (Phelan et al., 2006)), an association between blue color and organismal physiology has not been previously identified.

Notably, *C. xamachana* are exceptionally robust across variable environments; evidence suggests that their range is expanding globally across tropical coastal marine habitats (Bayha & Graham, 2014; Holland et al., 2004; Morandini et al., 2017; Stampar et al., 2020). *Cassiopea sp.* primarily inhabit shallow coastal ecosystems, where temperatures vary considerably among seasons and even within days (Aljbour et al., 2019; Kayal et al., 2018; Rowe et al., 2022). Studies show that *C. andromeda* are highly tolerant to stress and may even benefit from acute heat stress (Banha et al., 2020). In *C. xamachana,* this resilience appears to be independent of their symbiotic algae. *Cassiopea xamachana* that inhabit the Florida Keys are almost exclusively dominated by *Symbiodinium microadriaticum* (LaJeunesse, 2001). Although *S. microadriaticum* is generally considered to be thermally susceptible (Díaz-Almeyda et al., 2011; Iglesias-Prieto et al., 1992; Iglesias-Prieto & Trench, 1997; Warner et al., 1999), this group exhibits high levels of intraspecific variation in the response to temperature (Mansour et al., 2018). Understanding the underlying physiological mechanisms that enable *Cassiopea* to thrive under temperature stress may provide unique insights into the impact of global climate change on marine invertebrates.

In jellyfish, bell pulsation may play a role in oxygen exchange and temperature regulation, such that rates of bell pulsation typically increase with rising temperatures (M. N. Arai, 1996; Béziat & Kunzmann, 2022; Weller & Westneat, 2019), and decline rapidly as lethal levels are reached (Dillon, 1977). In *Cassiopea sp.*, bell pulsation rates have been suggested to reflect metabolic rates (Fitt et al., 2021; Mangum et al., 1972; McClendon, 1917). Thus, this can be used as an observable, whole-organism, ecologically-relevant phenotype for assessing organismal response during acute heat stress.

Here, we investigate how environmental history, appendage color, and temperature acclimation influence *C. xamachana* medusae responses to heat stress. Similarly, as in other cnidarians, we find that *C. xamachana* responds to acute heat stress by increasing bell pulsation rates until ∼37 ℃, at which point pulsation rates slow until lethality (38 – 41 ℃). Daily temperature ranges that varied between collection sites had no effect on rates of bell pulsation or lethal temperature. However, we present data showing evidence that *C. xamachana* can acclimate to increased temperatures and that survival is associated with the presence of blue appendages. This work is the first to demonstrate a link between color phenotype and resilience to thermal stress in *Cassiopea* and highlights a novel role for chromoproteins in thermal physiology.

## Methods

### Collection sites and environmental data

For all experiments, *C. xamachana* medusae were collected by hand from two sites in Key Largo, FL, USA. The “Atlantic” site (25.086669, -80.453382) is near-shore and receives water flow from the Atlantic Ocean. The “Bay” site (25.079426, -80.453009) is located off docks leading into the Florida Bay (Supplemental Figure 1). Water temperature was measured at collection sites using pendant HOBO® data loggers (MX2201) were deployed from March 12 to September 9, 2022 at a depth of 1.5 m to reflect the depth at which *Cassiopea sp.* are regularly observed. Daily mean temperatures and ranges (maximum - minimum) were calculated using 15-minute interval data (Figure 1A,B). Statistical differences between mean daily temperature and daily temperature range were determined using t-tests in R (v. 4.3.0; R Core Development Team 2017).

**Figure 1:**
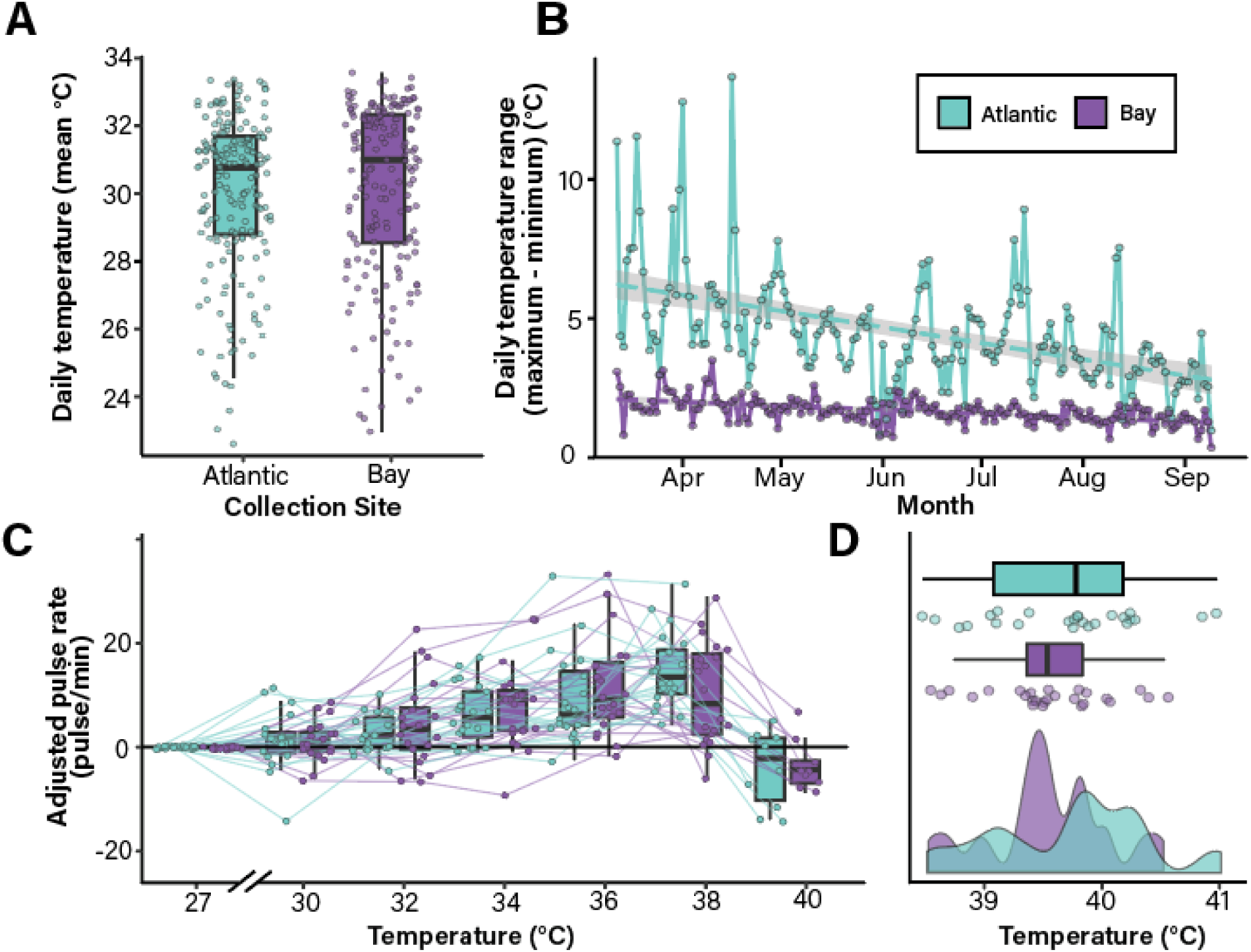
Environmental history does not affect *C. xamachana* bell pulsation rate or lethal temperature during acute heat stress. **A,B. *Cassiopea xamachana* that inhabit the Atlantic site experience greater environmental variation than those in the Bay site.** Daily mean temperatures (A) and temperature ranges (B) are shown for the Atlantic (teal) and Bay (purple) collection sites. Statistical analyses indicate that the two locations have similar daily mean temperatures (p_t-test_ = 0.378) but significantly variation in daily temperature (daily temperature range; p_t-test_ < 0.0001). **C,D. *Cassiopea xamachana* collected from the two sites do not differ in their response to an acute lethal heat stress.** Each point represents data from a single individual. Data are shown as adjusted pulse rate (the difference in bell pulsation rate (pulses per minute) between the baseline temperature (27 °C) and the temperature indicated). Lines connect measurements collected from individual medusa. A density plot of lethal temperatures is shown in D. Statistical analysis indicates that collection site does not influence lethal temperature during acute heat stress (p_lmer_ = 0.356).

### Experiment 1: Assessing the role of environmental variability on C. xamachana heat stress physiology

In May 2021, approximately 50 *C. xamachana* medusae (28 – 95 mm) were collected from each of the two sites (Supplemental Figure 1) and transported to Auburn University where they were acclimated for two weeks in a closed-sump aquarium system under ambient conditions (26 ℃ ± 1 ℃; 36 ppt salinity). Animals were maintained on a 12:12 light:dark cycle and fed *Artemia* three times per week. Morphological characteristics (bell diameter, sex, and appendage morphology) were recorded, and animals were re-acclimated for one week to compensate for handling stress.

Acute heat stress trials were performed such that 10 animals were monitored simultaneously and trials were repeated five times with different individuals (10 animals x 5 blocks = 50 total animals measured). Within each block, five individuals from each collection site were chosen at random. The evening before each trial, individual medusae were placed into 500 mL containers with mesh sides in the experimental tank (water conditions were identical to acclimation tank). Water temperature was controlled using two submersible rod heaters (Finnex Deluxe Titanium Tube Heater, 800W) connected to a digital temperature controller (SunTHIN WiFi Digital Temperature Controller 1250W, 10A). Experimental tank temperatures were monitored using pendant HOBO® data loggers (MX2201) recording temperatures at 15-minute intervals. Heat stress trials were performed at the same time each day to reduce potential variability associated with photoperiod (Nath et al., 2017). Water temperature was increased 1°C/hr; 30 minutes after each temperature increase, 1-minute videos were recorded in triplicate using Blink security cameras (Amazon) suspended above the experimental tank. Trials were complete when all medusae reached lethality, defined here as either 0.5 x baseline pulsation rate or < 10 pulses/minute. This criterion corresponded with tissue disintegration and was often accompanied by a separation of arms from bell tissue (Toullec et al., 2024). Dead animals were immediately removed from the experimental tank. Lethal temperatures and bell diameters at time of death were recorded for each individual.

### Experiment 2: Testing the role of temperature acclimation on C. xamachana heat stress physiology

In May 2022, approximately 50 *C. xamachana* medusae of each color morph (blue and uncolored/brown) were collected from the Atlantic site (Supplemental Figure 1) and transported to Auburn University where they were randomly placed into two sump systems, each with 4 individual tanks, and maintained as described above. After a five-day acclimation period, morphological characteristics (bell diameter, sex, and appendage morphology) were recorded. Individuals were photographed at a standard height (0.5 m) using a Canon Rebel X camera with a size and color standard. Tissue samples were collected from arm tips to measure symbiont density (stored at 4°C) and isolate genomic DNA (flash-frozen in liquid nitrogen and stored at - 80°C). Animals were acclimated for three days to compensate for handling stress.

The temperature in one of the two sump systems was increased 2 °C per day and then maintained at 33 °C (± 1 ℃) for 30 days (N=43 these animals are referred to as “elevated”). Individuals in the remaining tanks (N=42) were maintained in ambient conditions (26℃ ± 1℃). Temperatures were monitored using pendant HOBO® data loggers (MX2201). After the acclimation period, bell diameters were measured, and arm tissue was sampled to quantify symbiont density (stored at 4°C).

Acute heat stress trials were performed on medusae three days after post-acclimation measurements. Trials were performed as described in Experiment 1, with the additional parameters that 1) animals of each color morph were evenly distributed among the blocks; and 2). ‘elevated’ animals were transferred to the experimental tank when the temperature matched that of their acclimation tank (33 ℃).

### Bell pulsation analysis

Rates of bell pulsation were quantified using the videos captured during acute heat stress experiments. Videos were viewed once per individual within the camera view. Pulsation rate was calculated as the number of pulses per minute. To account for variation in pulsation rates among individuals, rates for each temperature were adjusted based on the baseline rate for each individual. For the 2021 trials and the ambient medusae from 2022, baseline rates were calculated before the temperature was changed in the experimental tank. For the elevated medusae in the 2022 trials, baseline rates were calculated in the experimental tank at 32 °C.

To test the effects of collection site or acclimation, appendage color, and temperature on bell pulsation rates, a generalized linear mixed-effect model (GLMM) with a Poisson link-log function was conducted in R using the package *lme4* (v.1.1-7, Bates et al., 2015). In these designs, location/acclimation, appendage color, and temperature during heat stress were fixed effects, and individuals and the date of the experiment (block) were coded as random effects. To test the effects of location/acclimation, appendage color, and heat stress temperature on the change in bell pulsation rates from baseline rates, a linear mixed effect model (LMM) was conducted the R package *lme4* (v.1.1-7, Bates et al., 2015). A LMM was used to examine differences in lethal temperature between location/acclimation and appendage colors. In this design, location/acclimation and appendage color were fixed effects; the date of the heat stress trial (block) was a random effect.

For all analyses, assumptions of normality and homogeneity of variance were graphically assessed using histograms, residual plots, and quantile-quantile plots. When there was a significant main effect (p < 0.05), Tukey’s post hoc mean comparisons were conducted using the *multcomp* package (v.1.4-25, Hothorn et al., 2008).

### Quantifying medusa appendage color

The amount of blue present in individual medusae was quantified by counting the number of pixels associated with blue using the R package *countcolors* (v.0.9.1, Weller & Westneat, 2019) and Adobe Illustrator. In brief, the background of each image was eliminated using Adobe Photoshop and replaced with white (manual editing was completed for each image to ensure the entire medusa was included). The *countcolors* package quantifies all pixels in an image according to their RGB (red, green, and blue channels; Weller & Westneat, 2019). The package was run using the following specifications: ignore white (set as background; RGB (1, 1, 1), total number of pixels will only include those that are not white), and identify pixels with an RGB value (the average quantified from all blue jellies in FIJI (Schindelin et al., 2012)) of (0.33, 0.30, 0.25) with an extended radius of 0.1; the *countcolors* package exports a new image with the pixels within the range ‘masked’ in magenta where the given pixel matches that of the parameters. Because the quantifiable color “brown” is similar to blue (blue: 0.25, 0.23, 0.21; brown: 0.52, 0.41, 0.23), images were checked in Adobe Illustrator to ensure brown appendages were not included. After any final modification, images were returned to the *countcolor* package and the total number of pixels and the number of magenta (RGB: (1, 0, 1)) pixels were quantified. A Pearson correlation was conducted in R to assess the relationship between lethal temperature and amount of blue coloration.

### Quantifying symbiont density

Medusa arm tip pieces were added to pre-weighed 1.5 ml centrifuge tubes and weighed to determine wet mass. Frozen tissue was homogenized in 500-700 μL of artificial seawater (Instant Ocean) using a disposable pestle and an electric homogenizer. To separate host tissue from the *Symbiodiniaceae* cells, homogenized samples were centrifuged at 800 x g for 10 minutes. Algal pellets were washed in 500 µl artificial seawater, vortexed to mix and re-homogenized as above. Samples were washed three times in artificial seawater (400 x g for 10 minutes) and resuspended in 150 µl Z-Fix (ANATECH). Symbionts were counted using an Improved Neubauer hemocytometer and normalized to the wet mass to calculate density. Eight counts were performed for each individual. A repeated measures ANOVA in R was used to examine symbiont density before and acclimation. In this design, acclimation treatment and appendage color were categorized as fixed effects; individual medusa ID was used as the repeated measure, and therefore the random effect.

### Verifying Cassiopea species

DNA from arm tissue was extracted using the E.Z.N.A. Tissue DNA Kit (Omega Bio-Tek) according to manufacturer’s instructions. DNA concentrations were quantified using the dsDNA Broad-Range Qubit assay (Invitrogen). The mitochondrial gene cytochrome c oxidase subunit I [COI] was amplified using custom-designed primers (Supplemental Table 1; e et al., 2012). Thermal cycling conditions were 3 min at 95°C for initial denaturation, followed by 35 amplification cycles (denaturation at 95°C for 35 s, annealing at 49°C for 40 s and extension at 72°C for 50 s) and a final extension for 7 min at 72°C. Amplified products were cleaned (Zymo Research Genomic DNA Clean and Concentrator-10 kit) and quantified using the dsDNA Broad-Range Qubit assay (Invitrogen). Purified products were sequenced using Sanger chemistry with custom primer MM1 (Supplemental Table 1). COI sequences were aligned to publicly available sequences from *C. xamachana* (Genbank accession NC_016466.1(Ohdera et al., 2019)) and *C. andromeda* (OP503353.1 and OP503325.1; (Kayal et al., 2012)) using Geneious v. 2022.2.2.

### Identifying Symbiodiniaceae type using ITS2 amplicon sequencing

Medusae collected from the Bay (n = 6) and Atlantic sites (n= 5) were used to characterize symbiont communities. Tissues samples were collected from arms, stored in ethanol and used for DNA extractions as described. ITS-2 regions were amplified using GoTaq MasterMix (Promega), and *Symbiodiniaceae*-specific primers (final concentration 400 nM) with adaptors for Illumina sequencing (Supplemental Table 1). Reactions were amplified as follows: 95°C for 3 min, 25 cycles of 95°C for 30 sec, 55°C for 30 sec, 72°C for 30 sec, and 72°C for 5 min. Samples were purified with the Genomic DNA Clean and Concentrator-10 kit (Zymo Research) and sequenced on the Illumina MiSeq platform (150 bp paired-end reads; Georgia Genomics).

The R package dada2 (Callahan et al., 2016) was used to identify and quantify the *Symbiodiniaceae* taxa present in each sample. After removing chimeric sequences, valid amplicon sequence variants (ASVs) were identified and enumerated. To identify the proportion and identity of the symbiont community in each sample, we created a BLAST database containing annotated ITS2 sequences from the SymPortal framework (https://symportal.org).

## Results

### Animals used in this study are *C. xamanchana*

Given the morphological variation and potential for cryptic species among *Cassiopea sp*., (Muffett & Miglietta, 2023) the species identity of collected animals was confirmed. DNA isolated from medusae used for the 2022 acute heat stress experiment was used to amplify the COI gene for sequencing. All 30 sequences exhibited >99.6% sequence identity with the *Cassiopea xamachana COI* sequence ((Kayal et al., 2012); Supplemental Figure 2a). Further, 11 representative animals from the collection sites were selected for ITS-2 symbiont typing. These medusae were overwhelmingly dominated by *Symbiodinium A1* (Supplemental Figure 2b), which is consistent with studies showing strong host-symbiont fidelity between *C. xamachana* and *S. microadriaticum* (LaJeunesse, 2017). Together, these data strongly indicate that, despite the morphological variation among individuals, medusae used in these studies are *C. xamachana*.

### Daily temperature variation does not affect rates of bell pulsation or lethal temperatures during acute heat stress

To determine if environmental history affects how *C. xamachana* responds to heat stress, medusae were collected from two sites that vary significantly in daily temperature range (“Atlantic” and “Bay” sites; Supplemental Figure 1). Temperature data collected from the two sites revealed that, although these locations had similar average daily temperatures (30.1 °C in the Bay and 29.9 °C in the Atlantic; Figure 1A), the Atlantic site water temperatures were significantly more variable than those in the Bay site (*p_t-test_* < 0.001, Figure 1B). On average, water temperatures at the Atlantic site varied 4.55 °C per day compared to 1.73 °C at the Bay site. This variability may reflect changes in water depth due to tidal influx.

Animals from each site were exposed to an acute lethal heat stress (starting from ambient temperature and increasing 1 °C/hour until lethality). Bell pulsation rates were used to monitor organismal response. Notably, no significant differences were observed in the change in bell pulsation rates among animals collected from the two sites (Figure 1C; Supplemental Figure 3; Supplemental Table 2). The rates of bell pulsation increased ∼3.1 pulses per minute for each 1 °C increase in water temperature (*p_GLMM_* < 0.001, Supplemental Table 3) until 37 °C, at which point pulsation rates dropped dramatically just prior to mortality. Post-hoc analyses identified statistical differences in the change in pulsation rates at several temperatures relative to baseline (Supplemental Table 4). Additionally, no significant difference was observed in average lethal temperatures of *C. xamachana* collected from the Atlantic (39.81 °C), and Bay sites (39.52 °C; *p_LMM_* = 0.356, Figure 1D, Supplemental Table 5). Notably, despite surviving to 40 °C, we observed no evidence of symbiont loss (bleaching) in medusa from either collection site. Together, these data indicate that, despite the difference in daily temperature range and variability between collection sites, *C. xamachana* from both locations displayed similar physiological phenotypes in response to acute heat stress.

### Temperature acclimation influences bell pulsation and lethal temperature

To determine if *C. xamachana* are able to acclimate to elevated temperatures, medusae (N=100) were collected from the Atlantic site and maintained at either ambient (26 °C) or elevated (32 °C) temperature for 30 days (Supplemental Figure 4A). The different acclimation temperatures did not cause significant differences in bell diameter or symbiont density (Supplemental Figure 4B,C). In fact, medusae in both treatment conditions exhibited reduced bell diameters and increased symbiont density (Supplemental Figure 4B,C), which may reflect the transition to the artificial aquarium system. Following acclimation, 40 individuals (N_ambient_ = 20, N_elevated_ = 20) were randomly selected for heat stress trials.

Following the 30-day acclimation period and after the first group of animals was subjected to the experimental heat stress, an unexpected mechanical failure in the ambient tank (26 °C) caused a temperature decrease of 5 °C for 24 hours. The remaining individuals were thus treated as “cold-shocked”. To account for the possibility that this affected subsequent responses, shock status was added as a fixed effect for all models. Notably, no significant interaction of shock status was observed with any other treatment factors. We have therefore interpreted fixed effects (temperature, acclimation treatment, and appendage color) equally.

Animals were exposed to an acute heat stress starting from the acclimation temperature and increasing 1 °C/hour until lethality. As in the initial study, rates of bell pulsation steadily increased with temperature until animals approached lethality (Figure 2; Supplemental Figure 5). Acclimation temperature had a significant effect on the change of bell pulsation rates from the start of heat stress (*p_LMM_* = 0.008, Figure 2A, Supplemental Table 6). Notably, medusae acclimated in elevated conditions survived to higher temperatures compared to those acclimated at ambient temperatures. The average lethal temperature for *C. xamachana* acclimated at ambient temperature was 39.97 °C compared with animals acclimated at elevated temperatures (41.2 °C, *p_LMM_* < 0.0001; Figure 2B; Supplemental Table 8). This data show that *C. xamachana* can acclimate to a consistent period of elevated temperatures and that this acclimation is beneficial in the context of acute heat stress.

**Figure 2:**
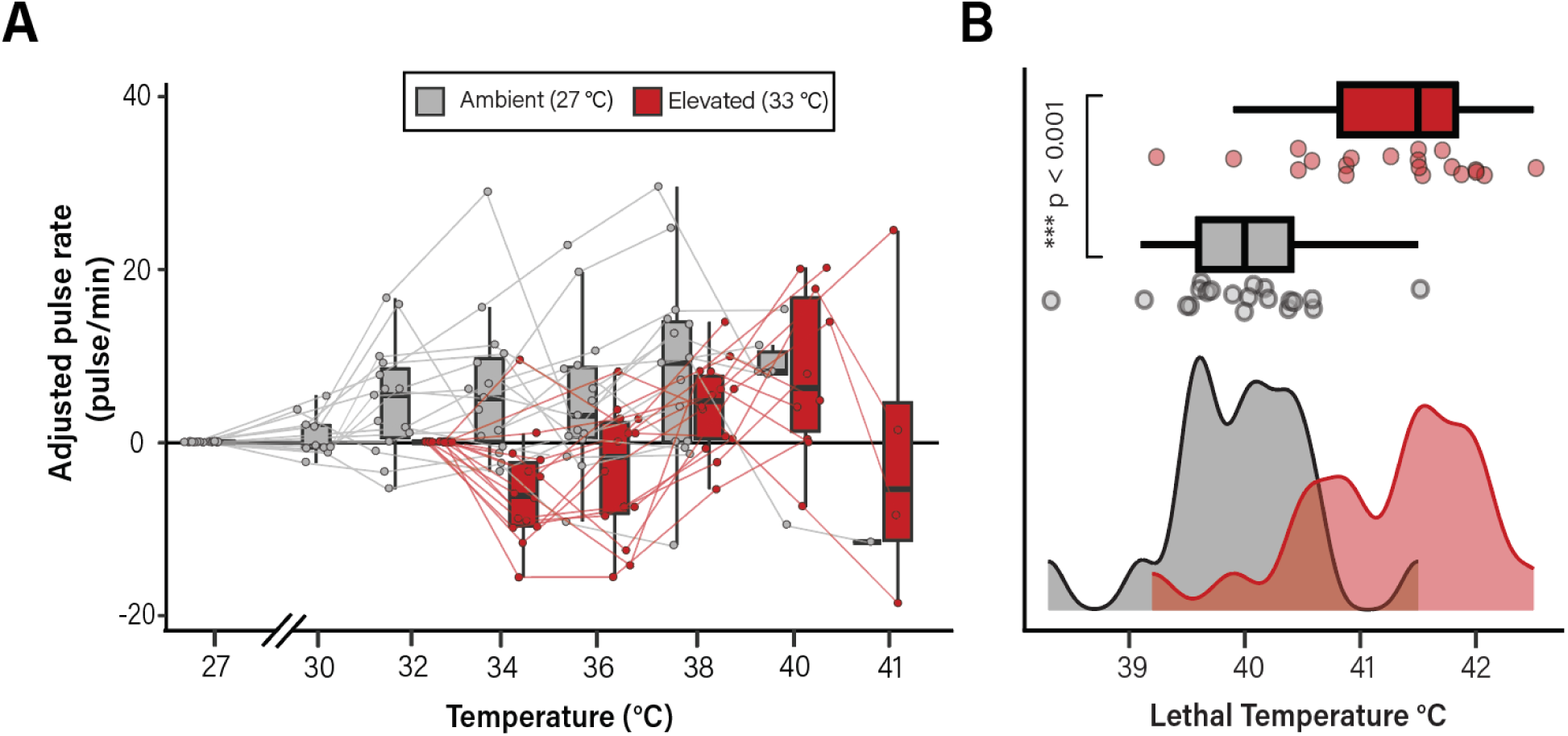
Temperature acclimation to significantly affects bell pulsation rate and lethality. Individual medusae were acclimated at either 27 ℃ (ambient; shown in red) or 32 ℃ (elevated; shown in black) and subsequently exposed to an acute lethal temperatures stress and monitored for rates of bell pulsation (**A**) and temperature at which death occurred (**B**). Data are shown as adjusted pulse rates such that, for each animal, the baseline rate (measured at either 27 ℃ for ambient or 32 ℃ for elevated individuals) was subtracted from the pulsation rate at the temperature indicated. Pulsation rates are shown as pulses per minute. Lines connect measurements collected from individual medusa. Data in B are shown as a histogram of lethal temperatures during the acute heat stress trial. A significant difference (*p_lmer_* < 0.001) was observed in the lethal temperature of *C. xamachana* acclimated at ambient vs. elevated conditions.

### Appendage color is associated with C. xamachana survival during acute heat stress

Given the lack of significant differences in medusae collected from two sites (Figure 1), we were interested in identifying other factors that might influence *C. xamachana* physiology during heat stress. We therefore investigated if this phenotype is associated with survival in increased temperatures. Using the data from both acute heat stress experiments, *C. xamachana* individuals were categorized as either “blue” or “brown” based on the predominant color of their oral appendages and data were assayed in the context of appendage color (Figure 3). Importantly, no relationship was observed between appendage color and either bell diameter or symbiont density (Supplemental Figure 6). The change in rate of bell pulsation during the acute heat stress experiments was not significantly different between medusae with blue appendages or brown appendages in either experimental year (2021 experiment, *p_LMM_* = 0.699, Figure 3A; 2022 experiment, *p_LMM_* = 0.681, Figure 3B). However, in both experiments, medusae with blue appendages survived to significantly higher temperatures than those without (Figure 3C; 2021 experiment, *p_LMM_* =0.003, Supplemental Table 5; 2022 experiment, *p_LMM_* < 0.007, Supplemental Table 8). This effect was also observed in medusae collected from both sites in 2021, such that individuals with blue appendages from the Atlantic site survived to significantly higher temperatures than those from the Bay site (*p_LMM_* = 0.0013; Figure 3D; Supplemental Table 9). Appendage color was also significantly associated with survival among medusae subjected to different temperature acclimation regimes (*p_LMER_* < 0.0001; Figure 3D).

**Figure 3:**
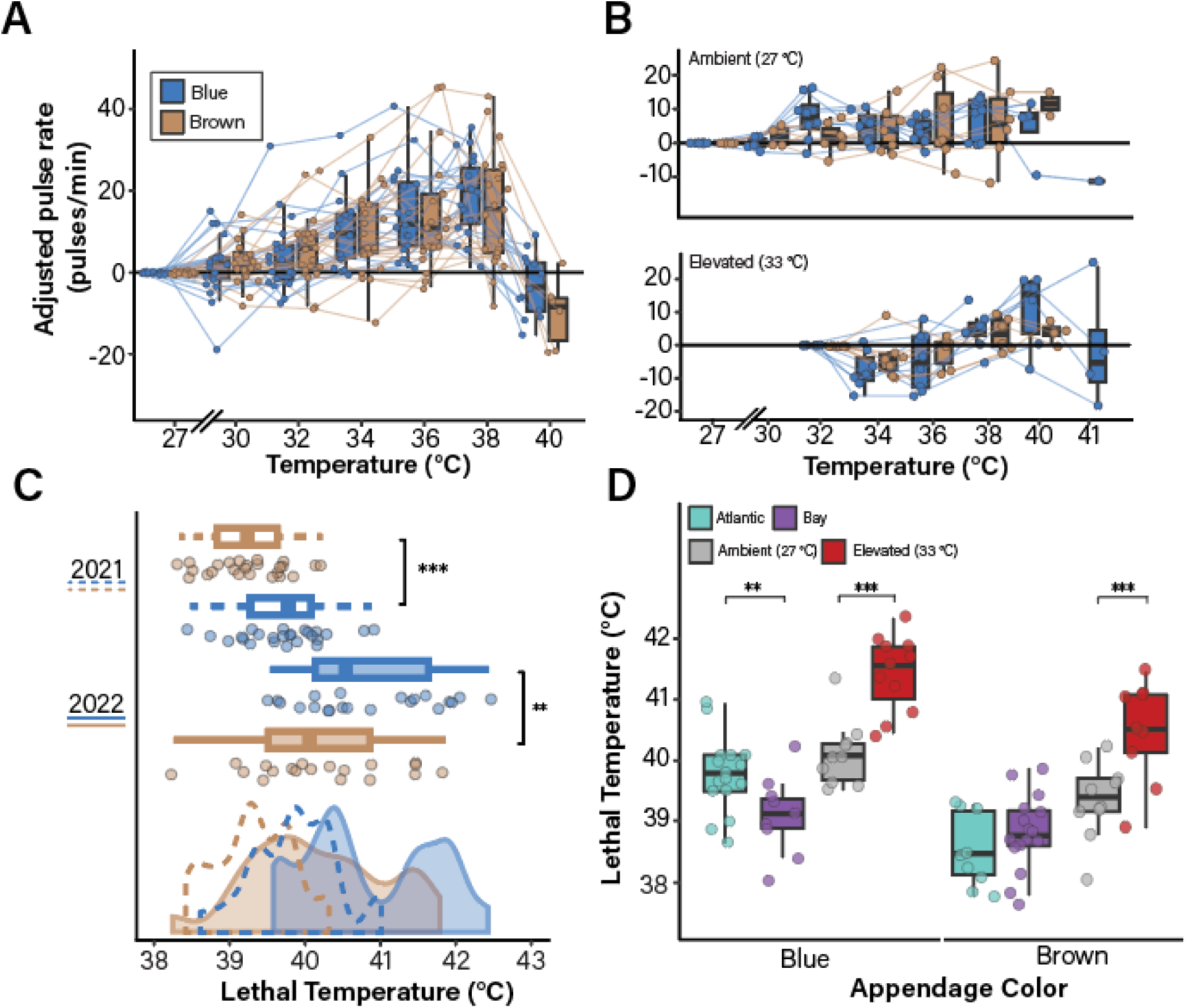
The presence of blue appendages is associated with *C. xamachana* survival during acute heat stress. Data from the experiments described in Figure 1 and Figure 2 were re-analyzed in the context of oral appendage color. Animals were classified as either “blue” or “brown” based on the predominant appendage color. No significant difference was observed in the adjusted pulsation rats of blue vs. brown animals in either experiment (2021 collection site data is shown in **A**, *p_lmer_* = 0.44; 2022 acclimation data is shown in **B**, *p_lmer_* = 0.94). Lines connect individual medusa between measurement points. Black dots indicate the final pulsation rate measured before lethality. **(C,D) Appendage color affects lethal temperature during acute heat stress**. Lethal temperatures are shown for the medusae collected in 2021 (C; *p_lmer_* = 0.0013) and 2022 (D; *p_lmer_* = 0.004). The effect of appendage color on lethal temperature in the context of collection location (*p_lmer_* = 0.0007, atlantic = teal, bay = purple) and acclimation treatment (*p_lmer_* = 0.004, gray = ambient, red = elevated) are shown.

To further investigate if there is a relationship between the amount of blue coloration and lethal temperature during an acute heat stress, appendage morphology was quantified as the number of pixels within the average RGB of blue appendages. A significant, positive correlation was observed between the percentage of blue coverage within a medusa and lethal temperature (R^2^ = 0.48; *p* = 0.010). Analysis of the acclimation treatment data revealed significant positive correlations between percent blue coloration and lethal temperature in medusae acclimated in either temperature treatment, although this effect was stronger in ambient conditions (R² = 0.34, *p* = 0.01), relative to elevated (R² = 0.15, *p* = 0.15, Figure 4).

**Figure 4:**
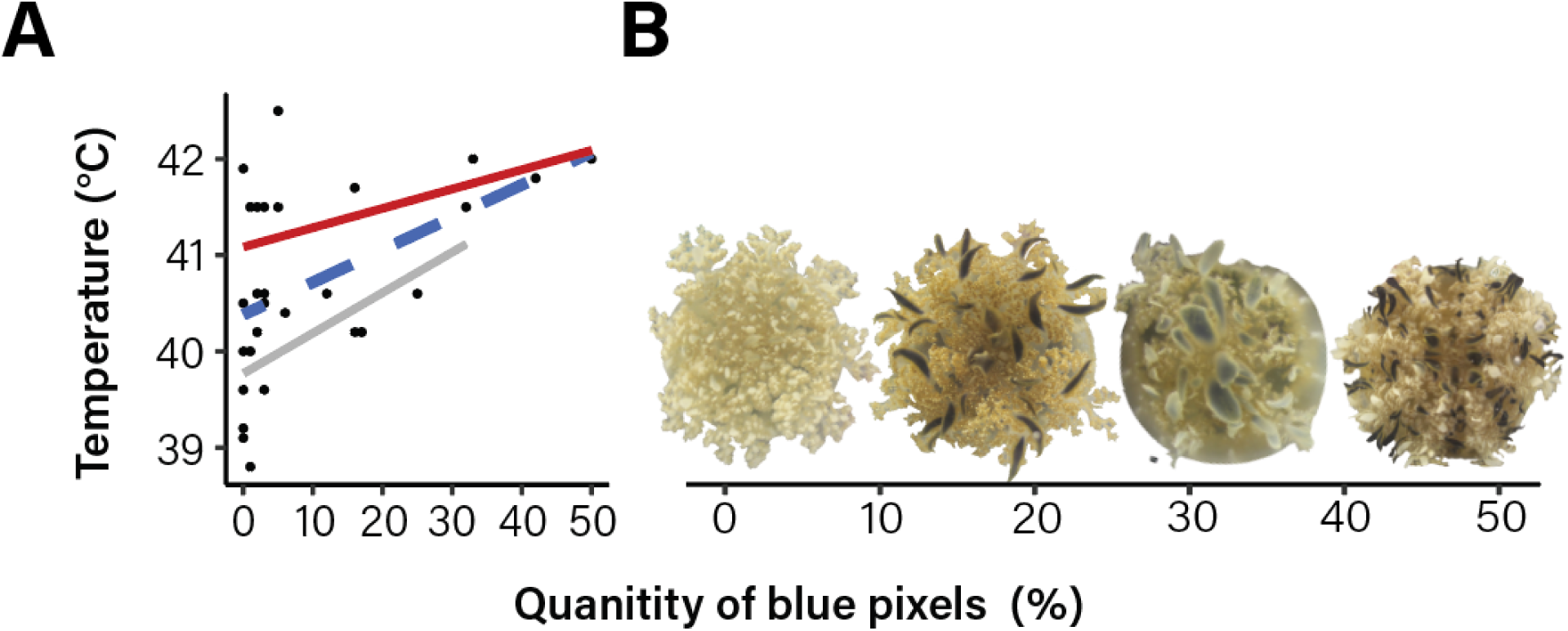
The amount of blue oral appendage correlates with survival during acute heat stress. **(A)** Animals acclimated at both ambient (gray; R² = 0.34, *p* = 0.01) and elevated (red; R² = 0.34, *p* = 0.01) temperatures benefit from the presence of blue appendages. Points represent individual medusa, lines represent the Pearson correlation between lethal temperature and quantity of blue pixels for all data points (blue dotted), ambient (grey) and elevated (red) acclimated medusa. (B) Representative images of medusa with pixel quantity ranging from 0 (left) to 50 (right) are shown.

## Discussion

Here, we present results that highlight how temperature regimes and coloration shape the ability of upside-down jellyfish (*Cassiopea xamachana*) to respond to thermal stress, revealing mechanisms underlying the remarkable resilience of this species. We find that animals with different environmental history, specifically differences in daily temperature range, do not exhibit variation in their thermal stress responses. However, short-term acclimation and the presence of blue appendages can improve organismal survival. Ultimately, these findings highlight a novel link between blue coloration and resilience in this marine invertebrate.

### Temperature influences the rate of bell pulsation in Cassiopea xamanchana

Among marine cnidarians, *Cassiopea* appears to be uniquely capable of thriving in a range of environmental conditions (Purcell, 2012; Toullec et al., 2023). Specifically, *C. andromeda* exhibits exceptional tolerance to repeated, intense (26 – 34 °C) heat stress episodes (Banha et al., 2020; Béziat & Kunzmann, 2022). Here, using rates of bell pulsation as a measure of organismal stress, we find that *C. xamachana* maintain homeostatic bell pulsation over a broad, but ecologically-relevant, temperature range (26 – 34 °C). This contrasts with several species of tropical cnidarians in which bell pulsation rates change dramatically beyond a narrow temperature range (Béziat & Kunzmann, 2022; Gatz et al., 1973). In *Cassiopea sp*., bell pulsations create water currents that facilitate food capture (Gohar & Eisawy, 1960; Hamlet et al., 2011; Klein et al., 2016) and oxygenate the surrounding water (Rowe et al., 2022). In the context of temperature stress, increased rates of bell pulsation may serve as a mechanism to enhance metabolic processes during stress (Fitt et al., 2021; Mangum et al., 1972; McClendon, 1917). This phenomenon has been observed in other marine invertebrates responding to acute temperature stresses (*e.g.*, crayfish; (Ern et al., 2015); gastropods; (Valles-Regino et al., 2022)). In *Cassiopea sp.*, Aljbour et al., (2019) found that antioxidant systems, such as SOD (superoxide dismutase) detoxifying activity (McCord & Fridovich, 1969), oxidative stress (OS) due to elevated formation of reactive oxygen species (ROS) (Regoli & Giuliani, 2014), and metabolic rate all correlated with rapid fluctuations in temperature (Aljbour et al., 2019; Fridovich, 1973; Lesser, 2006; McCord & Fridovich, 1969; Regoli & Giuliani, 2014). In stress conditions, such as increased temperature beyond physiological limits, high metabolic energy demand may increase reactive oxygen species (Boveris & Chance, 1973; Speakman & Selman, 2011). The ability to overcome oxidative stress could explain *Cassiopea sp.* resilience to thermal stress. Although *Cassiopea sp.* maintains a symbiotic relationship with algae similar to corals, the metabolic and physiological responses of the two do not necessarily mirror one another. For example, *Cassiopea sp*. have a unique breakdown of their symbiotic relationship. In this study, heat stress resulted in enhanced host catabolism coupled with reduced algal symbiont contribution, leading host bell diameter shrinkage “invisible bleaching (Toullec et al., 2024). Future work should aim to clarify the physiological and metabolic benefits or costs of increasing bell pulsation and how this contributes to the remarkable thermotolerance of *C. xamachana*.

### Environmental history does not shape how Cassiopea respond to acute heat stress

Theory by Lande, (2014) demonstrates that high environmental variability can result in higher phenotypic plasticity. This is supported by several empirical studies in marine invertebrates, demonstrating that organisms living in fluctuating environments (*e.g.,* drastic fluctuations in temperature or salinity) exhibit increased resistance to temperature stress associated with climate change (Burton et al., 2022; Godefroid et al., 2023; Kapsenberg & Cyronak, 2019; Rivest et al., 2017; Safaie et al., 2018). In contrast, here we demonstrate that *Cassiopea xamachana* collected from a site with high daily fluctuations in temperature exhibit the same physiological (bell pulsation rate) and survival (lethal temperature) response during heat stress than individuals collected from a site with stable daily temperature. Similar experiments in other cnidarian species have produced contradicting results. For example, temperature fluctuations on daily or tidal timescales were suggested to be sufficient to promote thermal tolerance of some corals via acclimation, while high temperatures were short enough to avoid lethality (Castillo & Helmuth, 2005; Oliver & Palumbi, 2011; Safaie et al., 2018). However, the effect of temperature variability on heat tolerance is likely species-specific (Putnam & Edmunds, 2011; Schoepf et al., 2022b; Voolstra et al., 2020).

Here we show that *C. xamanchana* are robust to thermal stress, and that this was not likely explained by inhabiting environments with high daily temperature range. Several potential factors may contribute to this result. First, the temperature fluctuations between the two sites may not be sufficiently different to elicit distinct physiological effects. Corals from habitats such as the back-reef pools in American Samoa experience daily temperature fluctuations up to 5.6 °C, even reaching daily extremes of >35 °C (Barshis et al., 2010); while the nearby, less-variable forereef has seasonal maximum daily temperature fluctuations of 1.8 °C (Barshis et al., 2018; Craig et al., 2001). These corals have phenotypic differences consistent with local adaptation of thermal performance (Palumbi et al., 2014). Similarly, either the temperature or the duration of daily maximum temperature in the Atlantic site may not be sufficient to induce increased thermal tolerance (either via acclimation or local adaptation).While our COI data revealed that all animals used in this study are *C. xamanchana,* (consistent with sequencing results from (Muffett & Miglietta, 2023)), our data is unable to resolve population-level differences or test for local adaptation. Thus, it is possible that *Cassiopea* sampled from both sites represent the same population and are thus robust to stable and fluctuating temperatures. These results are consistent with the overall observed robustness of *C. xamachana* relative to other marine (Aljbour et al., 2019; Banha et al., 2020; Béziat & Kunzmann, 2022; Fitt et al., 2021; McGill & Pomory, 2008; Medina et al., 2021; Ohdera et al., 2018; Rowe et al., 2022). In fact, *Cassiopea sp.* thrive in human disturbed environments (Stoner et al., 2011, 2016) and are expanding their habitat range. Our results may therefore reflect the underlying thermal tolerance of *Cassiopea* rather than environmental conditions of specific collection sites.

### Acclimation allows Cassiopea to survive elevated temperatures

As oceans warm for more prolonged periods due to global change, marine organisms must adjust their physiology to tolerate greater extremes (Coumou & Rahmstorf, 2012; Lopez et al., 2018; Morley et al., 2019). Short-term temperature stress can lead to subsequent increases in thermal tolerance in cnidarians (Baker et al., 2008; Kenkel & Matz, 2016; Kirk et al., 2018). The work presented here indicates that acclimation to a warmer environment may promote survival under acute heat stress and thus may prepare *C. xamachana* medusae for future warming conditions. Given the rapid heating of shallow water environments, *Cassiopea* may have an advantage if previous warming allows them to acclimate; it is possible that the previously encountered temperature ‘extreme’ in our acclimation experiment could have buffered the effect of the acute heat stress, thereby allowing them to survive to higher lethal temperatures (i.e., stress memory; Crisp et al., 2016; Li et al., 2020).

These results, while novel for *Cassiopea xamachana.*, are not uncommon among marine ectotherms. In natural experiments, corals of the Great Barrier Reef pre-exposed to thermal stress exhibited protection from subsequent heat waves (Ainsworth et al., 2016; DeMerlis et al., 2022). It is hypothesized then, that an organismal thermal threshold might be a plastic trait, in which some individuals are able to extend the upper thermal limit over time (Buckley & Huey, 2016; Leung et al., 2021). These observations are consistent with the laboratory-based experiments presented here and by other groups investigating cnidarian thermal tolerance (Ainsworth et al., 2016; Bay & Palumbi, 2015; Bellantuono, Granados-Cifuentes, et al., 2012; Bellantuono, Hoegh-Guldberg, et al., 2012; Middlebrook et al., 2008). *Cassiopea* appear to be resilient to temperature stress, and this could be the result of a hardening capacity, or the ability to increase thermotolerance due to experiencing a prior non-lethal temperature stress (Sgrò et al., 2010). The results presented here emphasize *Cassiopea’s* ability to acclimate to warming temperatures but may also predict their invasion potential and survival in more diverse environments. Some studies have already investigated *Cassiopea’s* invasion into more temperate environments (Rowe et al., 2022), however it is possible, due to their ability to acclimate, they may expand their range into warmer habitats.

### Appendage color is associated with better survival in warming environments

How temperature-dependent processes differ among color morphs could impact species survival and influence how polymorphisms evolve and/or persist (Thompson et al., 2023). In the present study, we observed differences in *C. xamachana* response to heat stress in medusae containing blue appendages compared to those lacking blue appendages. These results were consistent between two years of experimentation with individuals collected from different sites, demonstrating the strength of our hypothesis that pigmented appendages may benefit *C. xamachana* during extreme heating events. To the best of our knowledge, this is the first record of appendage color contributing to fitness traits in *Cassiopea xamachana*. While blue appendage color is common among some *Cassiopea* species, many color morphs are commonly observed, including red, blue, green, brown, white, and purple (Lampert et al., 2012). In many organisms, color is associated with camouflage, sexual selection, and social interactions (Cuthill et al., 2017), however these roles are unlikely in *Cassiopea*. Given their habitat (shallow, clear seawater) and ecology (epibenthic, with brown coloration mimicking the substrate), bright blue coloration makes individuals more obvious. *Cassiopea* brood their larvae and therefore do not undergo mate selection based on coloration as in birds and reptiles. One potential role for blue coloration is photoprotection from the UV damage that may occur in well-lit, shallow habitats, but mechanistic studies have not been performed (Blanquet & Phelan, 1987; Phelan et al., 2006).

It was previously hypothesized that the pigmented areas of the host acted as refuge for the symbiotic algae partner and that certain colors were correlated with specific algal strains. However, this was refuted by (Lampert et al., (2012) as well as the presented data, where color morphs did not have an association with a particular symbiont (Supplemental Figure 2) and were found in pigmented and non-pigmented host tissues. This is also consistent with results from a non-symbiotic, but blue jellyfish, *Rhizostoma pulmo,* which has a similar predicted protein associated with blue color (Bulina et al., 2004; Lawley et al., 2021). *Cassiopea sp*. vary in their pigmentation, appendage shape and number (Lampert et al., 2012; Rowe et al., 2022); we found that medusa with higher percent coverage of blue pigmentation were associated with higher lethal temperatures.

Trade-offs between fitness benefits and energetic costs of producing pigmentation may impact organismal physiology (Calsbeek et al., 2010), however these factors may not be relevant during acute heat stress in *Cassiopea*. In some organisms, selection pressures for coloration may result in variation in color traits (Zajitschek et al., 2012). However, in *Cassiopea*, blue appendages do not appear to influence whole-animal performance traits, such as bell pulsation, but may contribute to overall survival through alternative mechanisms (Husak & Fox, 2008; Irschick et al., 2008; Zajitschek et al., 2012). Blue color in many organisms can be achieved by combining a chromophore with an apoprotein (Bulina et al., 2004), which is not a common phenomenon due to the complex structure of blue pigments. It therefore must be exceptionally beneficial to produce and have blue pigmentation in *Cassiopea*. While there are several predicted proteins associated with blue color in other Rhizostomins, including *R. pulmo,* other pigment colors have not yet been explored. While blue appendage color is common among some *Cassiopea* species, many color morphs are commonly found together, yet only the blue pigment has been explored. Future work on *Cassiopea sp.* pigmentation should investigate the source of other colors and determine both the molecular and physical benefits, should they exist.

## Conclusions

In the present study, we assessed the impact of environmental history, temperature acclimation and appendage color on the rate of bell pulsation and survival during an acute heat stress in *Cassiopea* jellyfish. Overall, we demonstrate that *C. xamachana* medusae increase bell pulsation rates during an acute heat stress until temperatures reach ∼37 °C, at which point the rates sharply decline as the animals approach lethality. Surprisingly, we found that environmental history did not affect bell pulsation or survival, but pre-conditioning *Cassiopea* to warmer temperatures enables acclimation and survival to more extreme temperatures. Finally, our data show for the first time that the presence of blue appendages corresponds to survival of *Cassiopea* under warming climates. Together, these data hint at novel physiological mechanisms controlling the organismal response to thermal stress. Understanding the mechanisms that allow species to survive environmental stressors has become increasingly important as climate change continues to threaten marine taxa. This work contributes to our broader understanding of how phenotypic plasticity and color variability can shape organismal survival.

## Data Availability

Raw data files and analysis scripts are available at https://github.com/mem0294/Maloneyetal2024.

