## Supplemental Figures and Tables for "Blue appendages and temperature acclimation increase survival during acute heat stress in the upside-down jellyfish, *Cassiopea xamachana*"

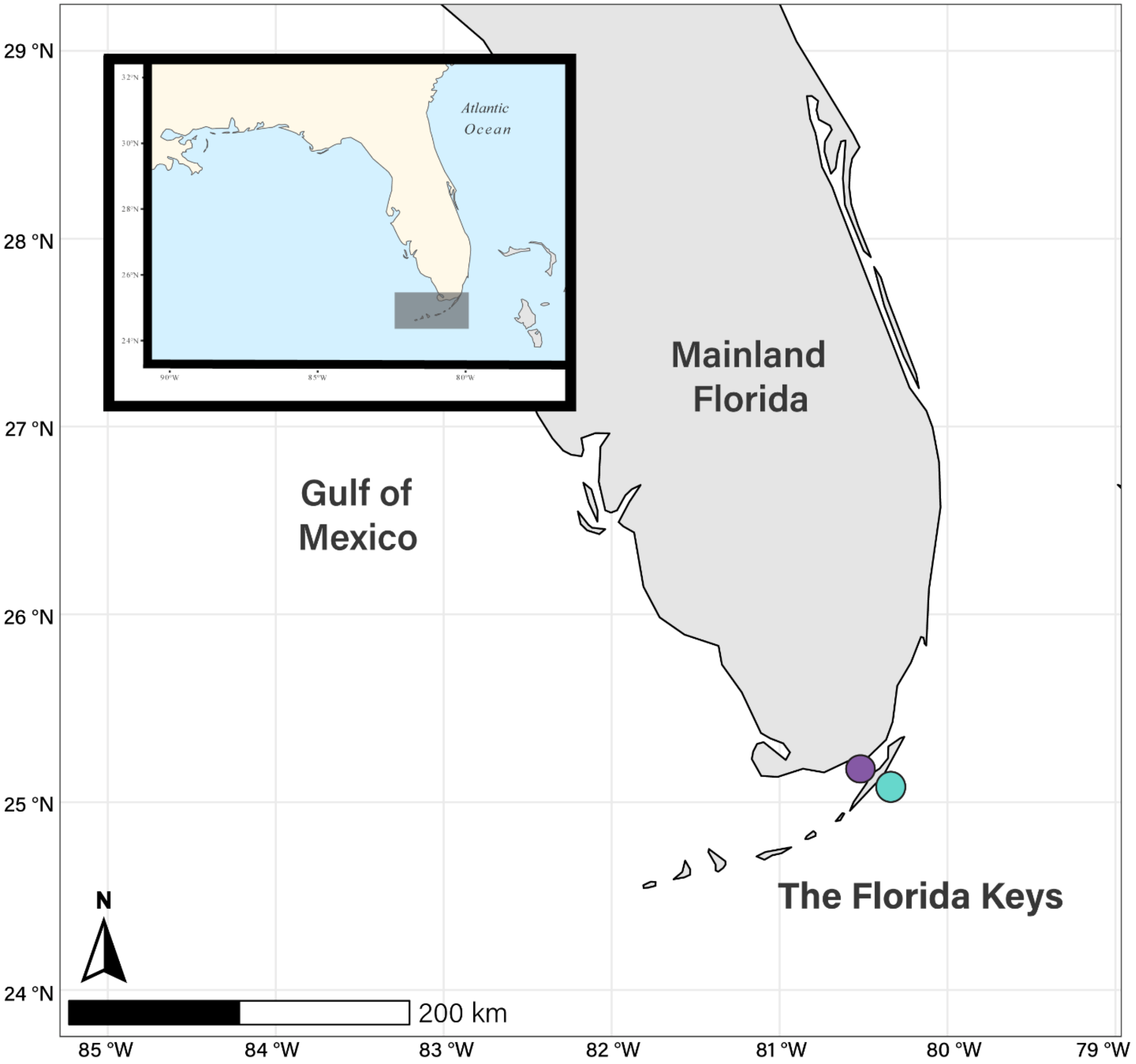


**Supplemental Figure 1:** ***Cassiopea xamachana*** **were collected from two distinct sites in the Florida Keys**. Approximate geographic locations for each collection site are shown. The Atlantic site (teal) is in a near-shore inlet surrounded by mangroves with a silt bottom. This site receives water flow from the Atlantic Ocean and is subject to tidal-based changes in water depth. The Bay site (purple) is located off docks that lead into the Florida Bay. Temperatures and depths are relatively consistent at this site (see Figure 1). In the 2021 study, approximately 50 individuals were collected from each site. In the 2022 study, 100 individuals were collected from the Atlantic site. All individuals were transported to Auburn University for long-term housing.

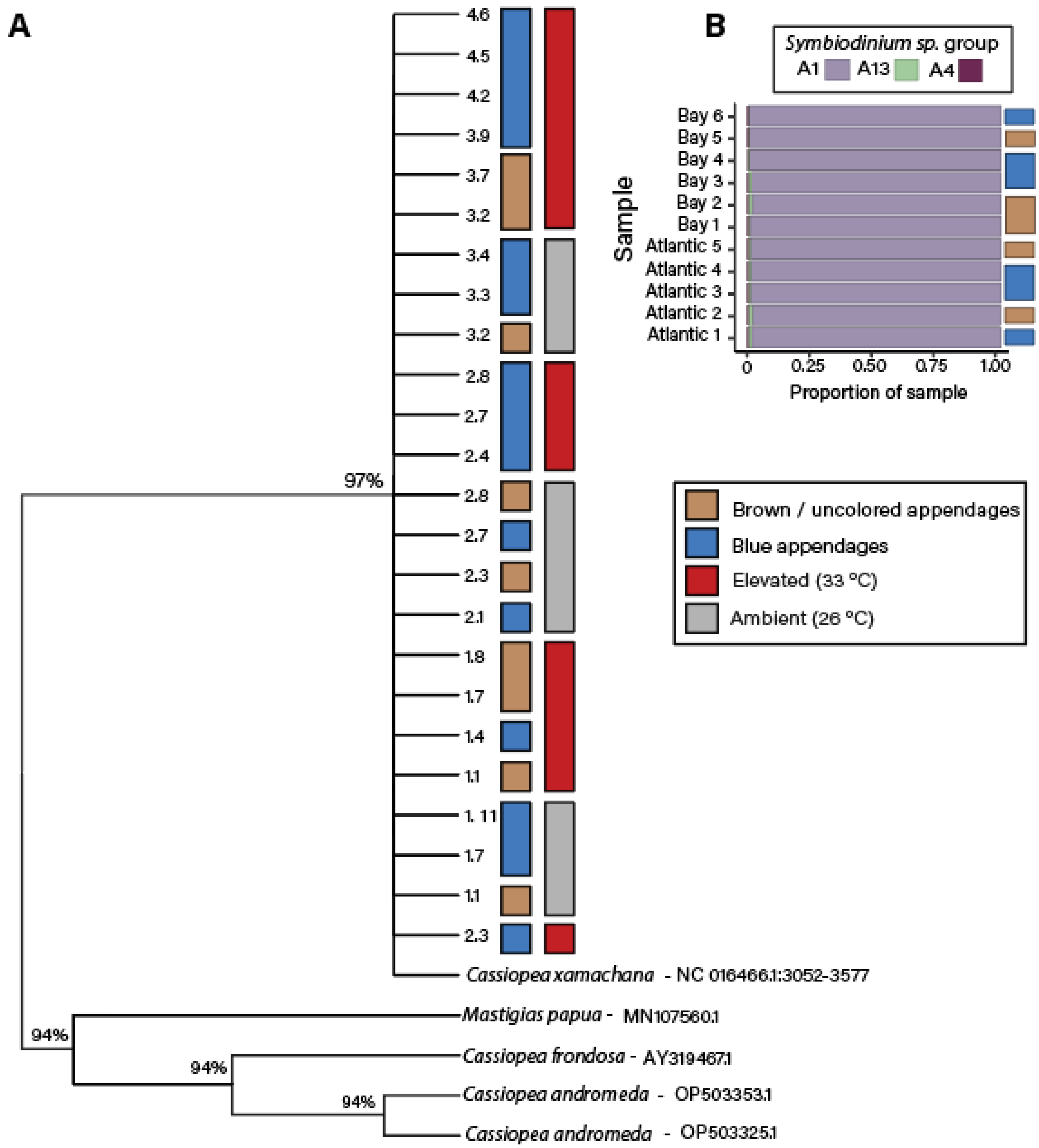


**Supplemental Figure 2: Medusae used in this study are *C. xamanchana*.** Sequencing the COI gene (A) and analysis of symbiont communities (B) indicate that the animals used here belong to the *C. xamanchana* clade, rather than other closely related species. COI gene sequences were aligned using the MUSCLE algorithm implemented in MEGA 10. Phylogenetic analysis was performed using maximum likelihood methods (Tamura et al., 2021). Bootstrap values based on 100 replicates are shown above each node. Experimental individuals are indicated with numbers (*e.g.*, 1.1). Appendage color (blue or brown) and acclimation temperatures (ambient [gray] or elevated [red]) are shown. NCBI accession numbers are shown for sequences from other *Cassiopea* species used in this analysis. (B) Normalized relative proportion of ITS2-type profiles from selected medusa from Bay and Atlantic ($N_{Atlantic} = 6$, $N_{Bay} = 5$). Barplot colors represent the symbiont community associated in each medusa, appendage color is indicated by blue or brown block to the right of the plot.

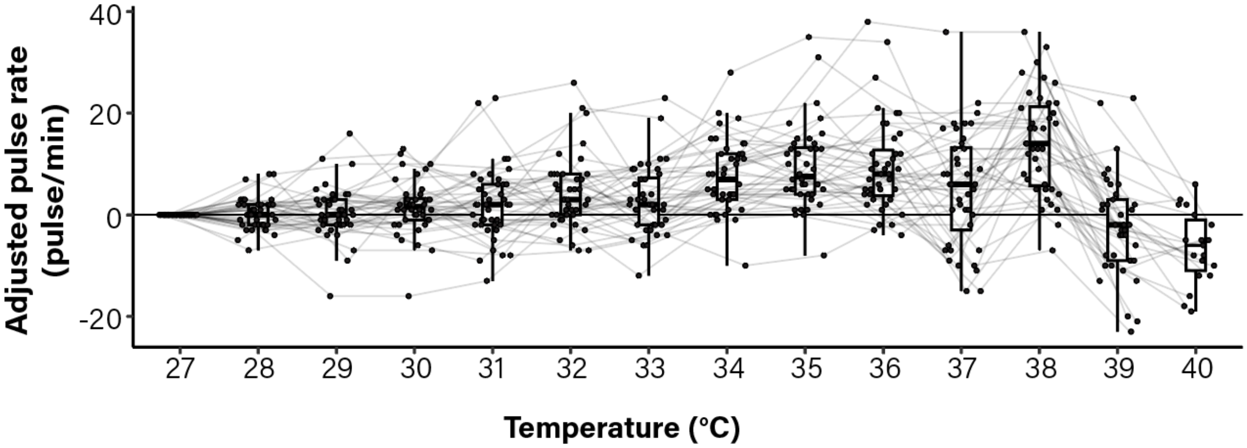


**Supplemental Figure 3:** Data are shown as adjusted pulse rate (the difference in bell pulsation rate (pulses per minute) between the baseline temperature (27 °C) and the temperature indicated). Measurements were collected after individuals were transferred to the experimental tank but prior to increasing the temperature. Lines connect measurement points from individual medusa.

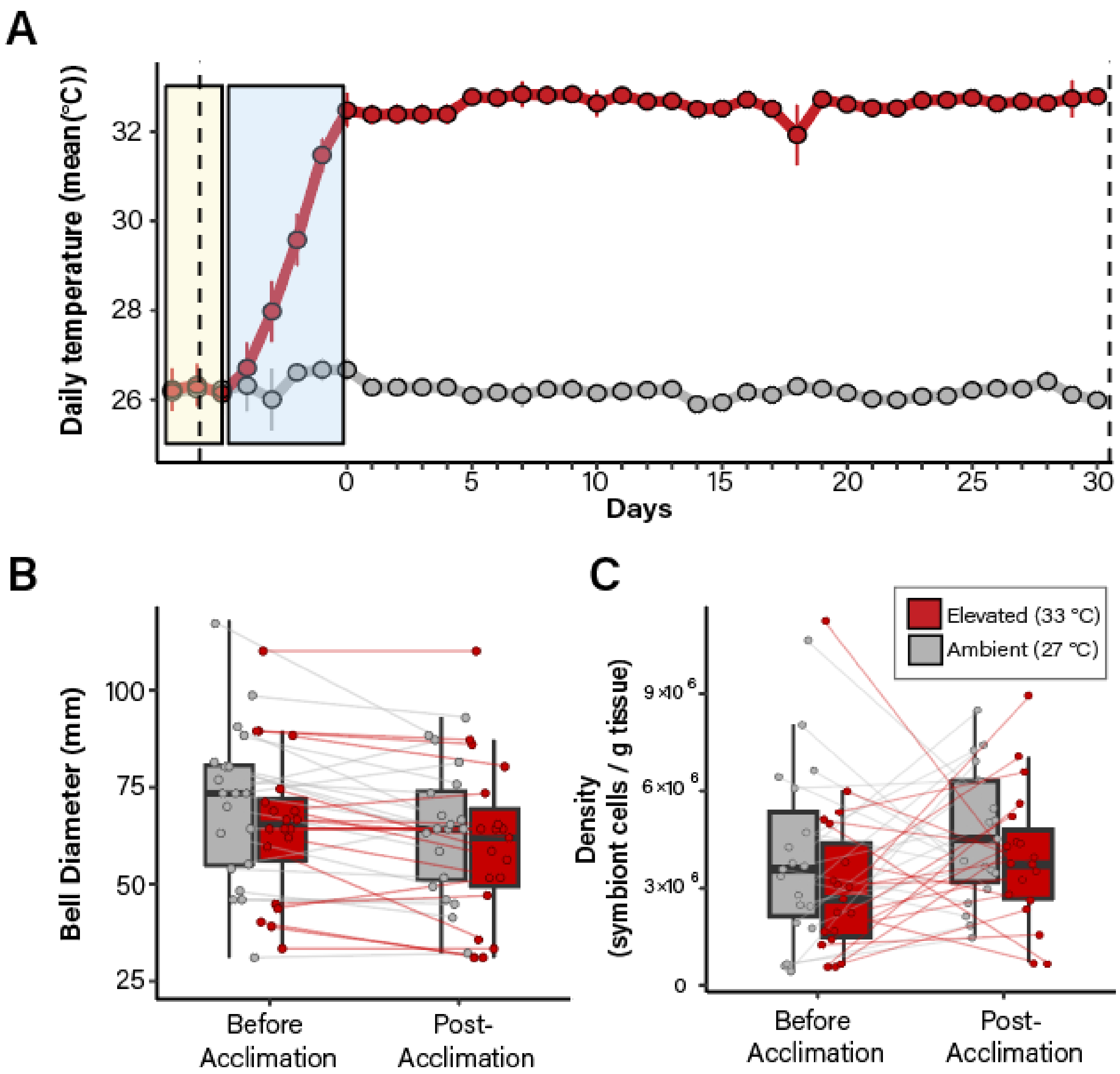


**Supplemental Figure 4:** (A) The experimental design used to assess temperature acclimation. Medusa were transported from collection sites and housed for three days (shaded yellow region) at Auburn University. Prior to increasing water temperature in the elevated temperature tanks, "before" measurements (bell diameter size, image, sex, color, symbiont and DNA sample) were collected (dashed line). Water temperature in the elevated tank system was increased 1-2 °C per day until it reached an average of 32 °C (blue shaded region; 5 days). Points represent the mean temperature in the systems (+/- s.d) Red dots represent the elevated condition, grey dots represent the ambient condition. "Post acclimation" measurements (size, image, and symbiont sample) were collected for each medusa after 30 days (dashed line). Heat stresses were performed after the 30-day period. (B,C) During the acclimation treatment, medusae did not exhibit a significant change in bell diameter (B) or symbiont density (C). Lines connect individual medusa between measurement points.

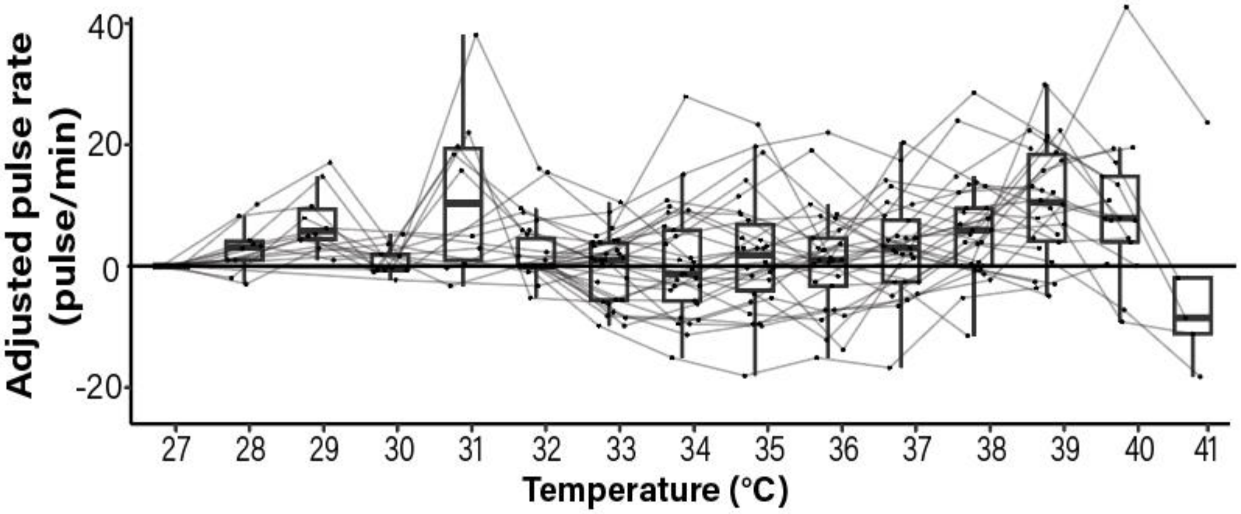


**Supplemental Figure 5: Change in pulsation rate during 2022 acute heat stress (post movement into experimental tank, but before temperature change).** Data are shown as adjusted pulse rate (the difference in bell pulsation rate (pulses per minute) between the baseline temperature (27 °C) and the temperature indicated). Measurements were collected after individuals were transferred to the experimental tank but prior to increasing the temperature. Lines connect measurement points from individual medusa.

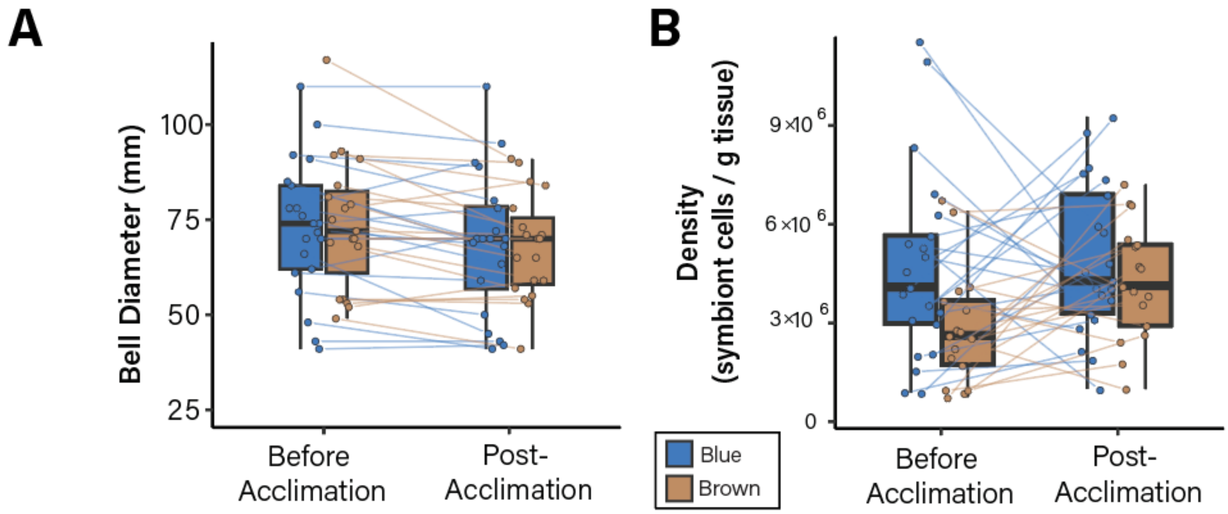


**Supplemental Figure 6:** During the acclimation treatment, medusae with differing appendage color did not exhibit a significant change in bell diameter (A) or symbiont density (B). Lines connect individual medusa between measurement points (before acclimation treatment vs. after).

**Supplemental Table 1: Primers used in this study.**

| Taxon | Target | Primer Name | Sequence (5 ′ - 3 ′) | Citation |
|---|---|---|---|---|
| *Symbiodiniaceae* | ITS-2 | SYM_VAR_5.8SII (+Illumina adaptor) | TCGTCGGCAGCGTCAGATGT GTATAAGAGACAGGAATTGC AGAACTCCGTGAACC | Hume et al. 2013, 2015, 2018 |
| | | SYM_VAR_REV (+Illumina adaptor) | GTCTCGTGGGCTCGGAGATG TGTATAAGAGACAGCGGGTT CWCTTGTYTGACTTCATGC | |
| *Cassiopea sp.* | COI | MM1F | AACGTCTTGGCATTCCTGCT | This study |
| | | MM1R | GCAGGTGGGGGAGATCCTAT | |

**Supplemental Table 2: Statistical results of a linear mixed effect model (LMM) for change in rate of *C. xamachana* bell pulsation from baseline (at 27 °C) at each temperature during heat stress[1].**

| Predictors | Estimates | CI | *p-value*[2] |
|---|---|---|---|
| 27 °C (Intercept) | -3.87 | -8.30 – 0.56 | 0.993 |
| 28 °C | 1.39 | -1.49 – 4.26 | 0.900 |
| 29 °C | 1.75 | -1.13 – 4.62 | 0.581 |
| 30 °C | 1.73 | -1.15 – 4.60 | 0.337 |
| 31 °C | 3.91 | 1.03 – 6.78 | **0.014** |
| 32 °C | 4.17 | 1.29 – 7.04 | **0.009** |
| 33 °C | 4.73 | 1.85 – 7.60 | **0.007** |
| 34 °C | 7.95 | 5.07 – 10.82 | **<0.001** |
| 35 °C | 9.99 | 7.11 – 12.86 | **<0.001** |
| 36 °C | 11.71 | 8.83 – 14.58 | **<0.001** |
| 37 °C | 15.29 | 12.41 – 18.16 | **<0.001** |
| 38 °C | 11.79 | 8.91 – 14.66 | **<0.001** |
| 39 °C | 1.88 | -1.06 – 4.81 | 0.224 |
| 40 °C | -4.93 | -8.51 – -1.34 | **0.047** |
| Location | 4.10 | 0.22 – 7.98 | 0.669 |
| Appendage Color | -1.90 | -5.90 – 2.10 | 0.699 |

[1] Random effect values are as follows: $\sigma_2$ = 50.33; $\tau_{00animal}$ = 12.87; $\tau_{00date}$= 7.13; $N_{animal}$ = 40; $N_{date}$ = 4; observations = 537; marginal $R^2$ / conditional $R^2$ = 0.234 / 0.629.
[2] Values in bold indicate significant differences between the intercept (temperature = 27°C, location = Bay, and Appendage color = blue) and the predictors listed.

**Supplemental Table 3: Statistical results from generalized linear mixed-effect model for rate of bell pulsation during heat stress of *Cassiopea xamachana*.**[1]

| Predictors | Incidence Rate Ratios | CI | *p-value*[2] |
|---|---|---|---|
| 27 °C (Intercept) | 21.26 | 17.65 – 25.62 | **<0.001** |
| 28 °C | 1.04 | 0.96 – 1.13 | 0.363 |
| 29 °C | 1.05 | 0.97 – 1.14 | 0.211 |
| 30 °C | 1.05 | 0.97 – 1.14 | 0.218 |
| 31 °C | 1.14 | 1.05 – 1.24 | **0.001** |
| 32 °C | 1.15 | 1.06 – 1.25 | **0.001** |
| 33 °C | 1.18 | 1.09 – 1.28 | **<0.001** |
| 34 °C | 1.31 | 1.21 – 1.42 | **<0.001** |
| 35 °C | 1.39 | 1.29 – 1.51 | **<0.001** |
| 36 °C | 1.46 | 1.36 – 1.58 | **<0.001** |
| 37 °C | 1.61 | 1.50 – 1.74 | **<0.001** |
| 38 °C | 1.47 | 1.36 – 1.59 | **<0.001** |
| 39 °C | 1.06 | 0.98 – 1.16 | 0.152 |
| 40 °C | 0.78 | 0.70 – 0.88 | **<0.001** |
| Location | 1.16 | 0.99 – 1.36 | 0.059 |
| Appendage Color | 1.01 | 0.86 – 1.20 | 0.865 |

[1] Random effect values are as follows: $\sigma_2$ = 0.04; $\tau_{00animal}$ = 0.08; $\tau 00_{date}$= 0.02; $N_{animal}$ = 50; $N_{date}$ = 5; observations = 720; marginal $R^2$ / conditional $R^2$ = 0.206 / 0.784.

[2] Values in bold indicate significant differences between the intercept (temperature = 27°C, location = Bay, and appendage color = blue) and the predictors listed.

**Supplemental Table 4: Tukey post-hoc results for the change in bell pulsation relative to baseline (2021 experiment).*[1]***

| Contrast | estimate | Std error | statistic | *p-value* (adj) | Significance |
|---|---|---|---|---|---|
| 34 - 27 | 7.6 | 1.524267 | 4.9860039 | 4.284E-05 | *** |
| 34 - 28 | 7.4 | 1.524267 | 4.8547933 | 0.0001335 | ** |
| 34 - 29 | 7.1 | 1.524267 | 4.6579773 | 0.0003301 | ** |
| 34 - 30 | 6.075 | 1.524267 | 3.9855229 | 0.0051811 | ** |
| 34 - 31 | 5.45 | 1.524267 | 3.5754896 | 0.024397 | * |
| 35 - 27 | 9.4 | 1.524267 | 6.1668996 | 1.132E-08 | *** |
| 35 - 28 | 9.2 | 1.524267 | 6.0356889 | 1.446E-07 | *** |
| 35 - 29 | 8.9 | 1.524267 | 5.838873 | 1.935E-07 | *** |
| 35 - 30 | 7.875 | 1.524267 | 5.1664185 | 1.19E-05 | *** |
| 35 - 31 | 7.25 | 1.524267 | 4.7563853 | 0.0001265 | ** |
| 35 - 32 | 5.25 | 1.524267 | 3.444279 | 0.0368435 | * |
| 35 - 33 | 6.55 | 1.524267 | 4.2971481 | 0.0011988 | ** |
| 36 - 27 | 9.625 | 1.524267 | 6.3145115 | 1.395E-08 | *** |
| 36 - 28 | 9.425 | 1.524267 | 6.1833009 | 1.22E-08 | *** |
| 36 - 29 | 9.125 | 1.524267 | 5.986485 | 1.073E-07 | *** |
| 36 - 30 | 8.1 | 1.524267 | 5.3140305 | 8.452E-06 | *** |
| 36 - 31 | 7.475 | 1.524267 | 4.9039973 | 7.499E-05 | *** |
| 36 - 32 | 5.475 | 1.524267 | 3.591891 | 0.0223233 | * |
| 36 - 33 | 6.775 | 1.524267 | 4.4447601 | 0.0007506 | ** |
| 37 - 27 | 5.275 | 1.524267 | 3.4606803 | 0.0347887 | * |
| 38 - 27 | 14.3 | 1.524267 | 9.38156 | 0 | *** |
| 38 - 28 | 14.1 | 1.524267 | 9.2503494 | 0 | *** |
| 38 - 29 | 13.8 | 1.524267 | 9.0535334 | 0 | *** |
| 38 - 30 | 12.775 | 1.524267 | 8.3810789 | 1.998E-15 | *** |
| 38 - 31 | 12.15 | 1.524267 | 7.9710457 | 2.753E-14 | *** |
| 38 - 32 | 10.15 | 1.524267 | 6.6589394 | 6.206E-09 | *** |
| 38 - 33 | 11.45 | 1.524267 | 7.5118085 | 4.658E-13 | *** |
| 38 - 34 | 6.7 | 1.524267 | 4.3955561 | 0.0009083 | ** |
| 38 - 37 | 9.025 | 1.524267 | 5.9208796 | 3.955E-07 | *** |
| 39 - 32 | -6.4319 | 1.55675 | -4.1316016 | 0.0025494 | ** |
| 39 - 34 | -9.8819 | 1.55675 | -6.3477564 | 5.86E-08 | *** |
| 39 - 35 | -11.682 | 1.55675 | -7.5040111 | 5.01E-13 | *** |
| 39 - 36 | -11.907 | 1.55675 | -7.6485429 | 1.796E-13 | *** |
| 39 - 37 | -7.5569 | 1.55675 | -4.8542607 | 7.508E-05 | *** |
| 39 - 38 | -16.582 | 1.55675 | -10.6515933 | 0 | *** |
| 40 - 27 | -6.5018 | 1.919652 | -3.3869878 | 0.0448649 | * |
| 40 - 28 | -6.7018 | 1.919652 | -3.4911734 | 0.0319613 | * |
| 40 - 29 | -7.0018 | 1.919652 | -3.6474517 | 0.0184991 | * |
| 40 - 30 | -8.0268 | 1.919652 | -4.1814028 | 0.0022704 | ** |
| 40 - 31 | -8.6518 | 1.919652 | -4.5069827 | 0.0007213 | ** |

| | | | | | |
|---|---|---|---|---|---|
| 40 - 32 | -10.652 | 1.919652 | -5.5488384 | 2.004E-06 | *** |
| 40 - 33 | -9.3518 | 1.919652 | -4.8716322 | 0.0001214 | ** |
| 40 - 34 | -14.102 | 1.919652 | -7.3460394 | 1.808E-12 | *** |
| 40 - 35 | -15.902 | 1.919652 | -8.2837095 | 1.776E-15 | *** |
| 40 - 36 | -16.127 | 1.919652 | -8.4009183 | 1.221E-15 | *** |
| 40 - 37 | -11.777 | 1.919652 | -6.1348822 | 2.322E-07 | *** |
| 40 - 38 | -20.802 | 1.919652 | -10.8362559 | 0 | *** |

[1] Only significant results are shown. Full results can be accessed at https://github.com/mem0294/Maloneyetal2024

**Supplemental Table 5: Statistical results from a linear mixed model testing the effect of collection site (location) and appendage color on lethal temperature of *C. xamachana* due to an acute heat stress.**

| Predictors | Estimates | CI | *p-value*[2] |
|---|---|---|---|
| (Intercept) | 39.99 | 39.75 – 40.23 | **<0.001** |
| Location | -0.15 | -0.46 – 0.17 | 0.356 |
| Appendage color | -0.49 | -0.81 – -0.17 | **0.003** |

[1] Random effect values are as follows: $\sigma_2$ = 0.02; $\tau_{00date}$= 7.13; $N_{animal}$ = 50; $N_{date}$ = 4; observations = 50; marginal $R^2$ / conditional $R^2$ = 0.215 / 0.943.
[2] Values in bold are significant.

**Supplemental Table 6: Statistical results from generalized linear mixed-effect model for rate of bell pulsation during heat stress (2022 experiment)[1].**

| Predictors | Incidence Rate Ratios | CI | *p-value*[2] |
|---|---|---|---|
| 27 °C (Intercept) | 19.80 | 15.76 – 24.87 | **<0.001** |
| 28 °C | 1.10 | 0.93 – 1.31 | 0.250 |
| 29 °C | 1.31 | 1.11 – 1.54 | **0.001** |
| 30 °C | 1.07 | 0.89 – 1.28 | 0.473 |
| 31 °C | 1.51 | 1.29 – 1.76 | **<0.001** |
| 32 °C | 1.24 | 1.08 – 1.42 | **0.002** |
| 33 °C | 1.10 | 0.96 – 1.26 | 0.180 |
| 34 °C | 1.24 | 1.08 – 1.42 | **0.002** |
| 35 °C | 1.23 | 1.07 – 1.40 | **0.003** |
| 36 °C | 1.15 | 1.01 – 1.33 | **0.040** |
| 37 °C | 1.28 | 1.12 – 1.47 | **<0.001** |
| 38 °C | 1.42 | 1.24 – 1.62 | **<0.001** |
| 39 °C | 1.64 | 1.43 – 1.87 | **<0.001** |
| 40 °C | 1.57 | 1.36 – 1.81 | **<0.001** |
| Acclimation | 0.97 | 0.82 – 1.16 | 0.768 |
| Appendage Color | 0.96 | 0.80 – 1.14 | 0.616 |

[1] Random effect values are as follows: $\sigma_2$ =0.4; $\tau_{00animal}$ = 0.06; $\tau_{00date}$= 0.01; N= 30; observations = 310; marginal $R^2$ / conditional $R^2$ =0.147 / 0.688.

[2] Values in bold indicate significant differences between the intercept (temperature = 27 °C, acclimation = ambient, and appendage color = blue) and the predictors listed.

**Supplemental Table 7: Statistical results from linear mixed-effect model (LMM) for change in rate of bell pulsation from baseline (27 °C or 32 °C)*[1]*.**

| **Predictors** | **Incidence Rate Ratios** | **CI** | ***p-value*[2]** |
|---|---|---|---|
| 28 °C | 0.34 | -4.41 – 5.10 | 0.887 |
| 29 °C | 1.89 | -3.87 – 7.65 | 0.519 |
| 30 °C | 6.42 | 0.66 – 12.18 | **0.029** |
| 31 °C | 1.19 | -4.56 – 6.95 | 0.684 |
| 32 °C | 10.92 | 5.16 – 16.68 | **<0.001** |
| 33 °C | 4.78 | 0.27 – 9.30 | **0.038** |
| 34 °C | 2.11 | -2.40 – 6.62 | 0.359 |
| 35 °C | 4.79 | 0.28 – 9.30 | **0.038** |
| 36 °C | 4.47 | -0.04 – 8.99 | 0.052 |
| 37 °C | 3.13 | -1.40 – 7.66 | 0.175 |
| 38 °C | 5.62 | 1.09 – 10.15 | **0.015** |
| 39 °C | 8.55 | 4.02 – 13.09 | **<0.001** |
| 40 °C | 12.98 | 8.33 – 17.62 | **<0.001** |
| Acclimation treatment | -4.84 | -8.42 – -1.25 | **0.008[3]** |
| Appendage color | -0.74 | -4.27 – 2.80 | 0.681 |

[1] Random effect values are as follows: $\sigma_2$ =50.51; $\tau00_{animal}$ = 18.72; $\tau00_{date}$= 1.58; N= 30; observations = 310; marginal $R^2$ / conditional $R^2$ =0.2603 / 0.431.
[2] Values in bold indicate significant differences between pulse rate at the temperature shown and the baseline temperature (27 °C or 32 °C depending on treatment).
[3] This value indicates a significant difference in the adjusted pulsation rates of animals subject to either acclimation temperature.

**Supplemental Table 8**: **Statistical results from a linear mixed model testing the effect of acclimation treatment and appendage color on *C. xamachana* lethal temperature during an acute heat stress.**

| ***Predictors*** | ***Estimates*** | ***CI*** | ***p-value***[2] |
|---|---|---|---|
| (Intercept) | 40.26 | 39.90 – 40.63 | **<0.001** |
| Acclimation treatment | 1.24 | 0.82 – 1.66 | **<0.001** |
| Appendage color | -0.60 | -1.02 – -0.18 | **0.007** |

[1] Random effect values are as follows: $\sigma_2$ =0.43; $\tau_{00block}$ = 0.00; $N_{block}$ = 4; $N_{animals}$ = 40; marginal $R^2$ / conditional $R^2$ =0.541 / NA.
[2] Values in bold indicate significant differences.

**Supplemental Table 9: Statistical results from a linear mixed effect model testing the effect of appendage color on lethal temperature (2021).**

| **Characteristic** | **Beta** | **95% CI** | ***p-value*** |
|---|---|---|---|
| Appendage color | -0.89 | -1.3, -0.47 | <0.001 |
| Appendage color * Location | 0.79 | 0.20, 1.4 | 0.010 |
